# Structural basis of ferroportin inhibition by minihepcidin PR73

**DOI:** 10.1101/2022.08.23.505007

**Authors:** Azaan Saalim Wilbon, Jiemin Shen, Piotr Ruchala, Ming Zhou, Yaping Pan

## Abstract

Ferroportin (Fpn) is the only known iron exporter in humans and is essential for maintaining iron homeostasis. Fpn activity is suppressed by hepcidin, an endogenous peptide hormone, which inhibits iron export and promotes endocytosis of Fpn. Hepcidin deficiency leads to hemochromatosis and iron-loading anemia. Previous studies have shown that small peptides that mimic the first few residues of hepcidin, i.e. minihepcidins, are more potent than hepcidin. However, the mechanism of enhanced inhibition by minihepcidins remains unclear. Here, we report the structure of human ferroportin in complex with a minihepcidin, PR73 that mimics the first 9 residues of hepcidin, at 2.7 Å overall resolution. The structure reveals novel interactions that were not present between Fpn and hepcidin. We validate PR73-Fpn interactions through binding and transport assays. These results provide insights into how minihepcidins increase inhibition potency and will guide future developments of Fpn inhibitors.

## Introduction

Ferroportin (Fpn) is a Fe^2+^/2H^+^ antiporter that is highly expressed in enterocytes, hepatocytes, and macrophages to export Fe^2+^ derived from either dietary intake or digestion of senescent red blood cells (Ganz, 2005; Pan et al., 2020; Vlasveld et al., 2019). Since Fpn is the only known iron exporter in mammals, its activity is essential for plasma iron homeostasis (Donovan et al., 2005). Ferroportin activity can be acutely suppressed by the peptide hormone hepcidin which directly inhibits iron transport activity and induces endocytosis and degradation of Fpn (Aschemeyer et al., 2018; Nemeth et al., 2004). Meanwhile, the expression of hepcidin is regulated via plasma iron levels (Babitt et al., 2007; Truksa et al., 2006). Mutations in Fpn that impair transport activity can cause ferroportin disease, leading to symptoms of iron deficiency anemia (Drakesmith et al., 2015; Pietrangelo, 2017). Elevated hepcidin levels can also lead to iron deficiency (Ginzburg, 2019). Hepcidin deficiency and hepcidin-resistant mutations in Fpn, on the other hand, lead to hereditary hemochromatosis and iron overload (Pietrangelo, 2017). Thus, the hepcidin-ferroportin axis must be tightly regulated to maintain serum iron levels.

Hepcidin is a peptide of 25 amino acids and is secreted by hepatocytes. Hepcidin binds to the extracellular side of Fpn (Billesbølle et al., 2020; Pan et al., 2020; Park et al., 2001). There are four pairs of intramolecular disulfide bridges in hepcidin (Hunter et al., 2002; Jordan et al., 2009) (**Figure 1a**), and mutational studies showed that the first 7 – 9 residues have a large impact on its ability to inhibit Fpn (Preza et al., 2011). The dissociation of the Fpn-hepcidin complex under reducing conditions led to the hypothesis that the disulfide bridge between Cys7 and Cys23 in hepcidin could be replaced with a disulfide bridge between Cys7 and Cys326 on Fpn (Fernandes et al., 2009), although this exchange was not observed in the structure Fpn bound to hepcidin (Billesbølle et al., 2020). These studies led to the development of a new class of potent Fpn inhibitors, termed minihepcidins, that are based on the first 7 – 9 amino acids of hepcidin. Some of the minihepcidins have drastically improved potency to Fpn and have been further optimized and tested as hepcidin replacements to treat patients with dysregulation of hepcidin (Ramos et al., 2012). However, it remains unknown how the minihepcidins bind to Fpn with a higher affinity than hepcidin.

**Figure 1.**
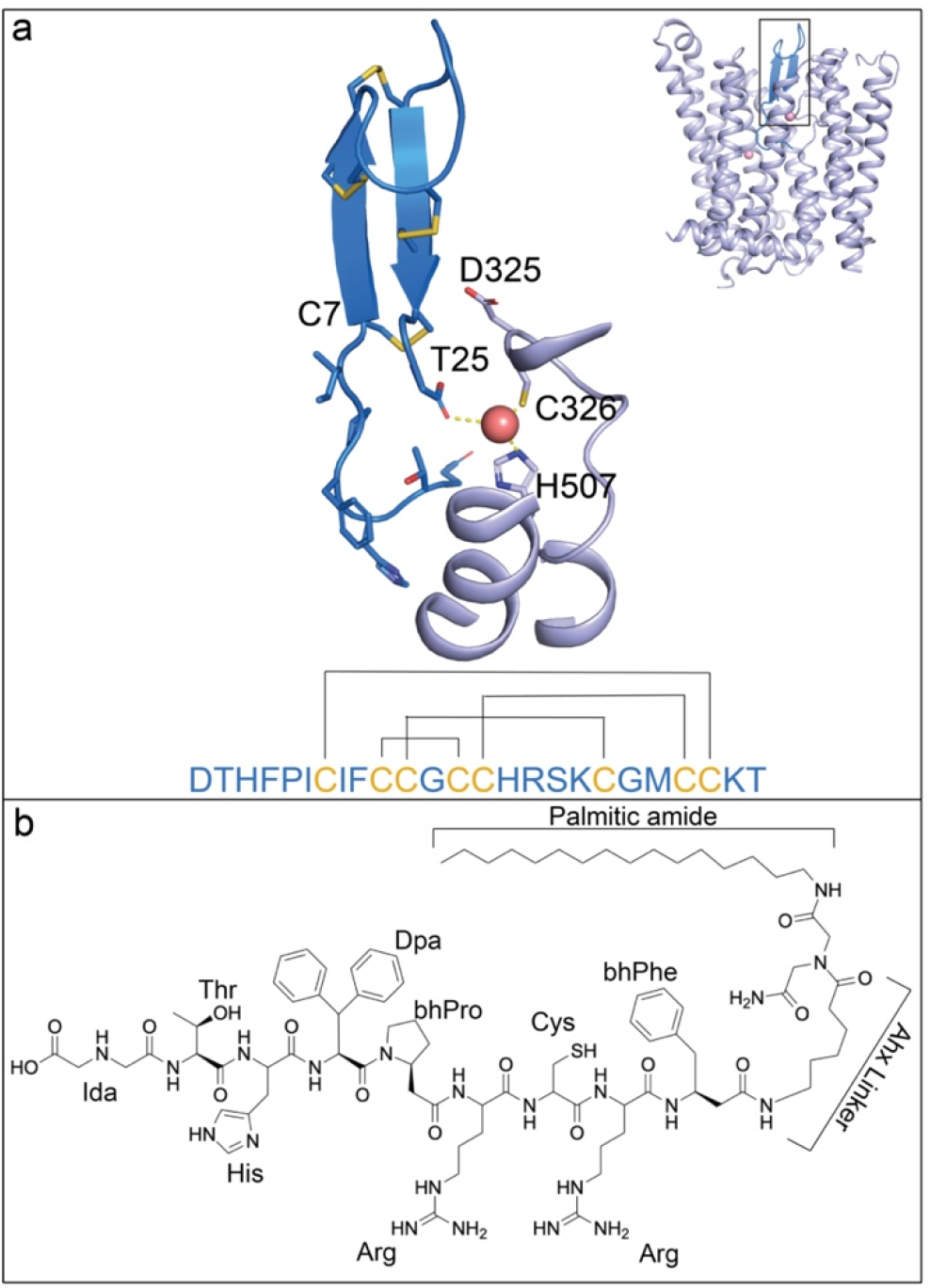
Structural comparison of hepcidin and PR73 molecules. **a**. Structure of hepcidin in complex with HsFpn (PDB: 6WBV). Hepcidin (marine), HsFpn (light blue), and Co^2+^ (light pink) are shown in the overall (inset) and zoomed-in views. Side chains of the first seven residues and all cysteines in hepcidin are shown as sticks, and so is the carboxyl group of the last residue. The hepcidin sequence is shown as single-letter codes at the bottom, and the disulfide bridges are marked by black lines. **b**. Chemical structure of PR73. Unnatural amino acid abbreviations are as follows: Ida = iminodiacetic acid, Dpa = diphenylalanine, bhPro = beta homo-proline, bhPhe9 = Beta homo-phenylalanine, Ahx = aminohexanoic linker.

Structures of Fpn in complex with hepcidin revealed how the two interact (Billesbølle et al., 2020; Pan et al., 2020). Hepcidin binds between the N-terminal domain (NTD) and C-terminal domain (CTD) of Fpn, and the first six residues of hepcidin interact with ferroportin. Hepcidin retains all four of its internal disulfide bonds, and its C-terminal carboxylate coordinates one metal ion at one of the binding sites, which is formed by Cys326 and His507 of Fpn (Billesbølle et al., 2020). Importantly, residue Cys326 is known to affect hepcidin binding and is associated with hemochromatosis (Fernandes et al., 2009; Ganz, 2006). These structures have presented insights into the binding mechanism of hepcidin to Fpn.

Minihepcidins are highly effective in inhibiting Fpn activity (Chua et al., 2015). PR73 is a minihepcidin and a highly potent inhibitor of Fpn (Fung et al., 2015) whose predecessor showed effectiveness in preventing iron overload (Ramos et al., 2012). PR73 mimics the first nine residues of hepcidin, including the cysteine in position seven that was hypothesized to form a disulfide bridge with residue Cys326 of Fpn (Fernandes et al., 2009) (**Figure 1b**). The last 16 residues of hepcidin have been replaced with an aminohexanoic linker and iminodiacetic palmitic amide (Ida(NHPal)) in PR73, eliminating the C-terminal carboxylate essential for the binding of hepcidin. Thus, it is not clear how PR73 inhibits Fpn with increased potency.

Here, we present the structures of human (*Homo sapiens*) Fpn (HsFpn) bound to PR73 and Co^2+^ at 2.7 Å and 3.0 Å, respectively. We used a fragment of antigen-binding (Fab) to facilitate structure determination by cryo-electron microscopy (cryo-EM). The structures show that PR73 preserves most of the interactions between hepcidin and Fpn. Although PR73 lacks the C-terminal carboxylate to coordinate the bound metal ion, it forms a disulfide bridge with HsFpn, and the palmitic amide of PR73 interacts with Gln194 on Fpn, which positions the acyl chain of PR73 in the hydrophobic core of the cell membrane surrounding Fpn.

## Results

### Structure of HsFpn in complex with 11F9 Fab

In a previous study, we reported the structure of the *Tarsius syrichta* ferroportin (TsFpn) in complex with an Fab from the mouse monoclonal antibody (11F9) (Pan et al., 2020). The Fab is crucial for structure determination because it increased the size of the particle and serves as a fiduciary marker for particle alignment. Since TsFpn is 92% identical to human Fpn and 15 of the 16 residues that form the epitope are identical in the two Fpns (**Figure 2—figure supplement 1a**), we tested the binding of 11F9 Fab to HsFpn. The Fab binds to HsFpn with a dissociation constant (*K*_*D*_) of 10.6 ± 3.2 nM (**Figure 2—figure supplement 1b**). In addition, we showed that HsFpn and the Fab can form a stable complex after reconstitution into lipid nanodiscs (**Figure 2—figure supplement 1c**). The 11F9 Fab is known to inhibit TsFpn activity (Pan et al., 2020), and we found that the Fab also inhibits Co^2+^ transport by HsFpn (**Figure 2— figure supplement 1d–e**).

**Figure 2.**
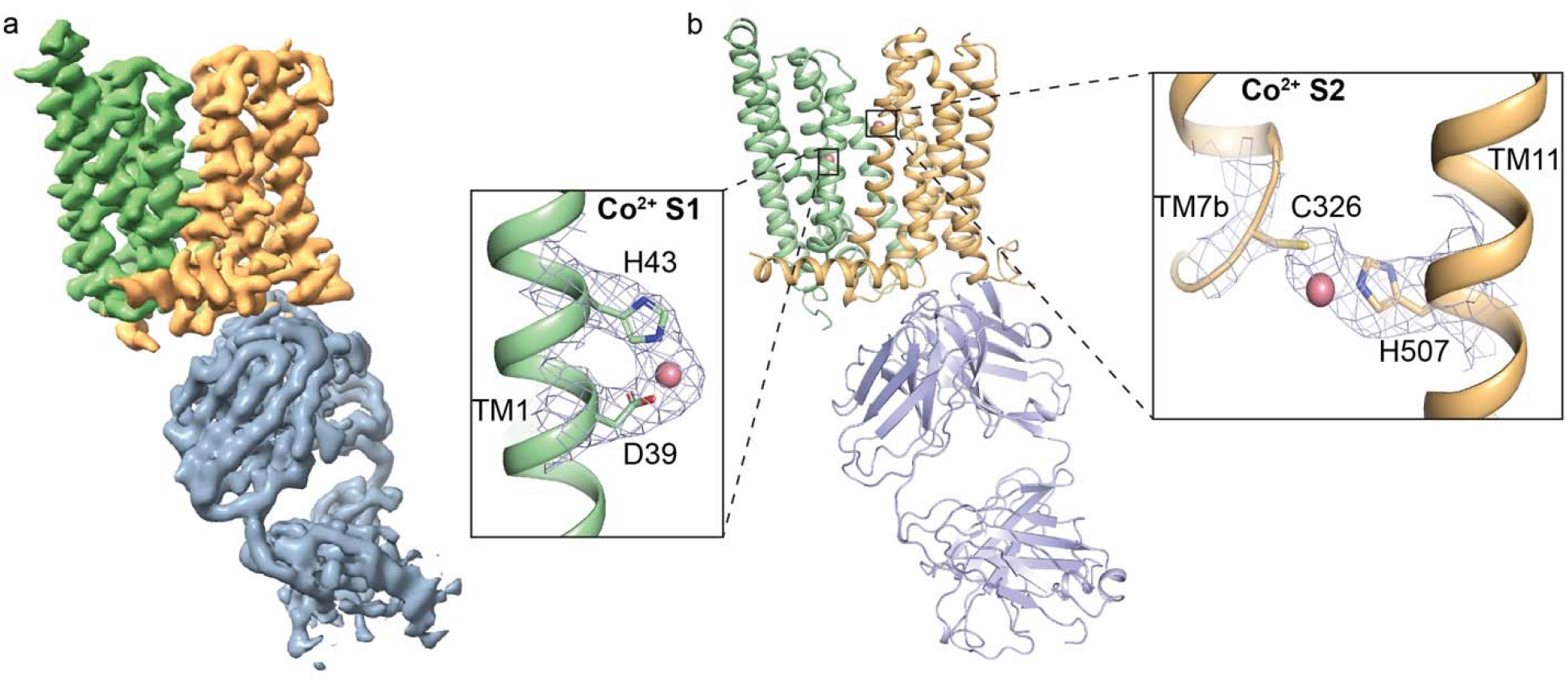
Structure of HsFpn in complex with Fab 11F9. **a**. Cryo-EM density map of HsFpn in complex with 11F9 in the presence of Co^2+^. Densities for the NTD, CTD, and Fab are colored in pale green, light orange, and slate grey, respectively and contoured at 8.5σ. **b**. Cartoon representation of HsFpn-11F9 complex. Ligand residues of the transition metal ion binding sites, S1 and S2, are highlighted and shown as sticks. Co^2+^ is rendered as a sphere (light pink). Densities for S1 and S2 are contoured at 4σ as blue mesh.

We reconstituted the HsFpn-11F9 complex into lipid nanodiscs and determined the structure of Co^2+^-bound HsFpn at an overall resolution of 3.01 Å by cryo-EM (**Figure 2a** and **Figure 2— figure supplement 2a–e**). The density map allows the model building of all transmembrane (TM) helices and most of the side chains. Residues 15 – 236, 284 – 396, and 451 – 557 are resolved and included in the final structure (**Figure 2—figure supplement 3**). Similar to previous structures of TsFpn and HsFpn, the current structure assumes an outward-facing conformation and aligns well with the previously reported human and monkey Fpn with a root-mean-squared distance (RMSD) of 0.57 Å and 0.58 Å respectively (**Figure 2—figure supplement 1a** and **f**). Two Co^2+^ binding sites are resolved, with the Site 1 (S1) composed of Asp39 and His43 and the Site 2 (S2) of Cys326 and His507 (**Figure 2b**). In addition, the Fab interacts with both the CTD and NTD of HsFpn, similar to the previously determined structures of TsFpn-11F9.

### Cryo-EM structure of Fpn bound to PR73

Next, we determined the structure of HsFpn-11F9 in nanodiscs in the presence of 1 mM of minihepcidin, PR73, to an overall resolution of 2.72 Å (**Figure 3a** and **Figure 3—figure supplement 1a–e**). The density map allows the modeling of residues 12 – 241, 283 – 416, and 449 – 558, which are included in the final structure (**Figure 3—figure supplement 2**). Residues 1 – 11, 242 – 282, 417 – 448, and 559 – 571 are not resolved. There are two lipid densities located near two amphipathic (AH) helices, one in the NTD and another in the CTD, on the intracellular side (**Figure 3—figure supplement 3**). A large non-protein density is present between the NTD and CTD towards the extracellular side, and was absent in previous maps of HsFpn when PR73 was not included. The density accommodates PR73 in an extended conformation (**Figure 3b–e**). The first 8 residues of PR73 occupy the cleft between the NTD and CTD of HsFpn, and the first 6 residues assume a similar conformation to that of hepcidin. A disulfide bridge is resolved between Cys7 of PR73 and Cys326 of HsFpn (**Figure 4b**), which confirms the previously proposed hypotheses (Fung et al., 2015; Preza et al., 2011). The binding of PR73 seems to push the NTD and CTD away from each other, which resembles what was observed in the hepcidin-bound HsFpn and TsFpn structures.

**Figure 3.**
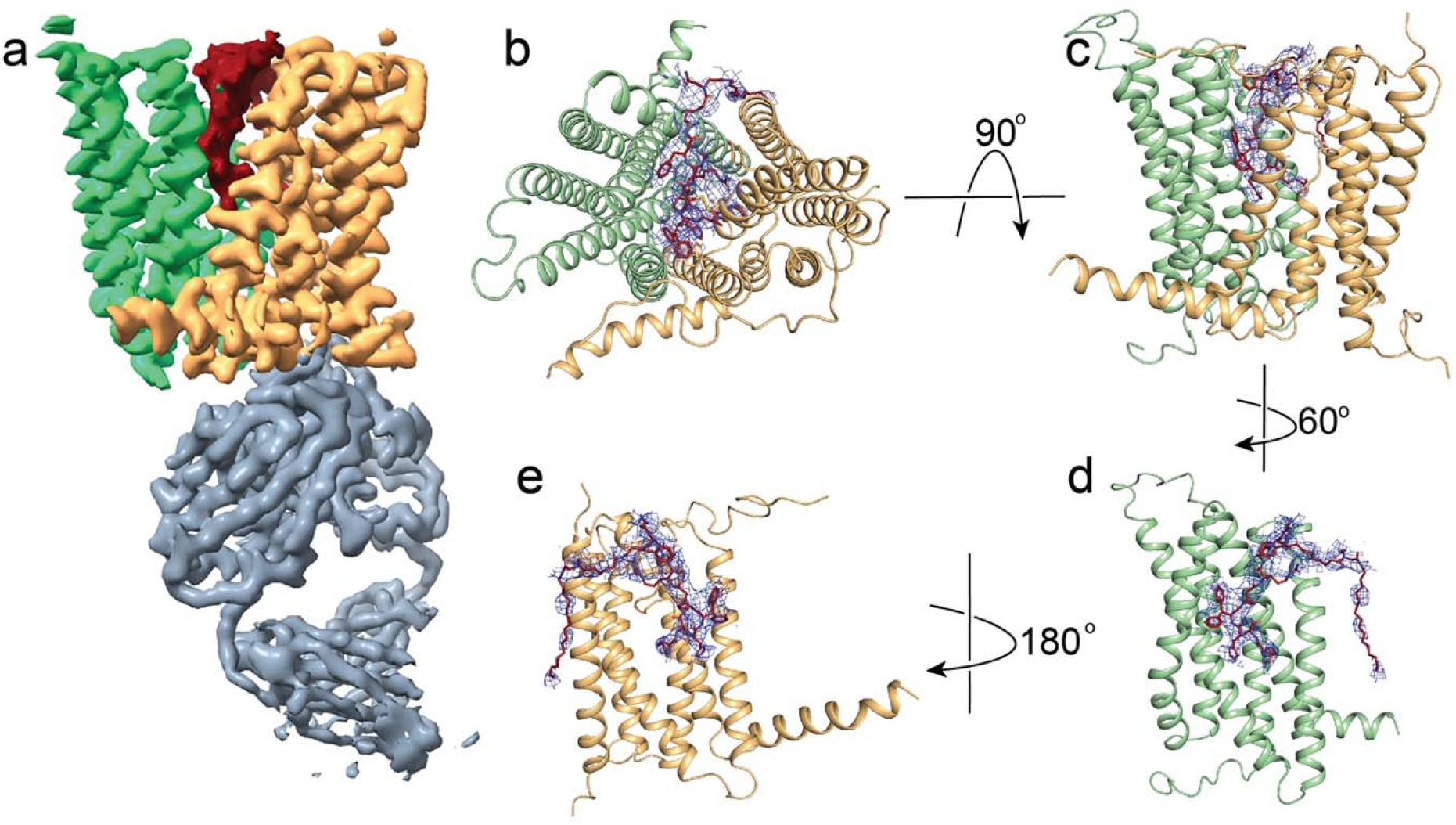
Structure of Fpn bound to PR73. **a**. Cryo-EM density map of HsFpn in complex with 11F9 Fab in the presence of PR73. Densities for the NTD, CTD, PR73, and Fab are colored in pale green, light orange, brick red, and slate grey respectively. HsFpn and the Fab are contoured at 8.5σ while the PR73 density is contoured at 4σ. Top-down **(b)** and side **(c)** views of the HsFpn-PR73 structure. HsFpn is shown as a cartoon, and PR73 is shown as sticks (brick red) with the density contoured at 4σ as blue mesh. Side views of HsFpn-PR73 with the CTD **(d)** or NTD **(e)** omitted.

**Figure 4.**
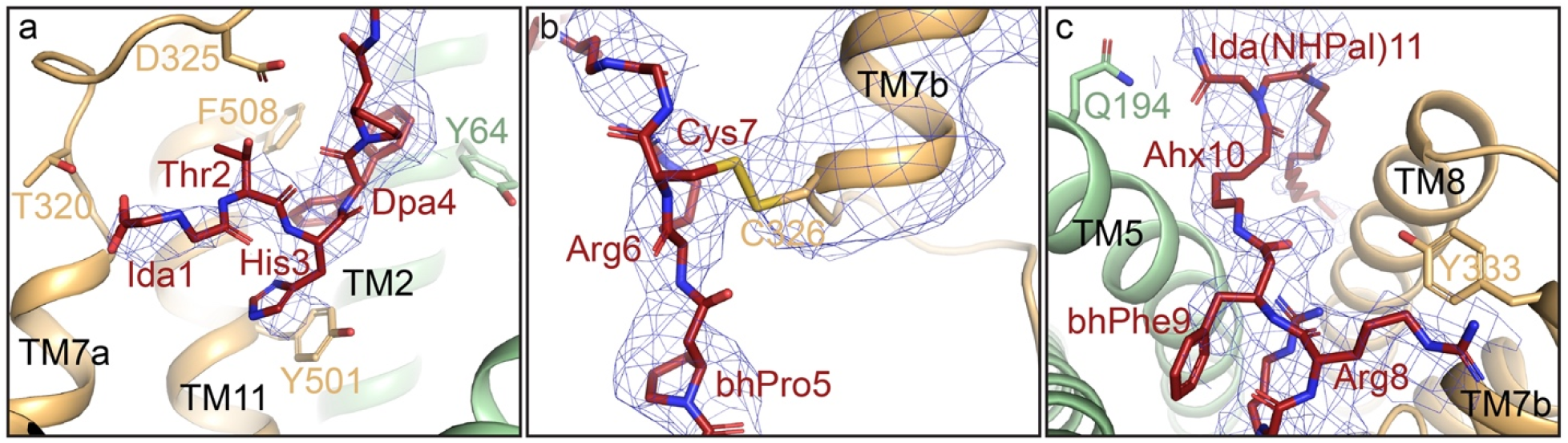
Interactions between PR73 and HsFpn. **a**. Zoomed-in view of the first four resides of PR73 peptide. The density of PR73 is contoured at 4.5σ as blue mesh. Residues of Fpn forming interactions with PR73 are shown as side-chain sticks. **b**. Zoomed-in view of residues 5 – 7 of PR73 displayed in the same representation as (**a**). The density of TM7b is contoured at 4.5σ as blue mesh. **c**. Zoomed-in view of the last four residues of PR73 shown in the same representation as in **a**.

The overall conformation of the PR73-bound HsFpn structure is similar to those of the hepcidin-bound HsFpn and TsFpn, with a C_α_ RMSD of 0.82 Å and 1.10 Å, respectively (**Figure 7**). When the CTD of the current structure is aligned to that of HsFpn without an inhibitor, the extracellular sides of the N-domain, especially of TM1, TM2, TM3, and TM4, are pushed away from the CTD (**Figure 7a–b**). The extracellular end of TM3 has the largest movement of ∼9 Å, while those of TM1, TM2, and TM4 have movements of 4 – 5 Å. In the CTD, the largest change is around S2. Here, TM7b has a rotation of ∼24° when compared to the hepcidin bound structure, likely induced by the disulfide bridge between Cys326 and Cys7 of PR73. This rotation is unique to the PR73 bound structure because in the hepcidin-bound structure Cys326 maintains its interaction with the bound ion, which in turn interacts with the C-terminal carboxylate of hepcidin.

Interactions between PR73 and HsFpn share common features with those of hepcidin and HsFpn, but there are several unique features as well. The first six residues of PR73 include native and modified versions of amino acids found in hepcidin. These residues interact with residues along TM7a and TM11 (**Figure 4a** and **Figure 4—supplement 1**), including residues Tyr501 and Phe508 which extend towards the hydrophobic region of PR73 between Thr2 and Arg6. These interactions are similar in the structure of hepcidin-HsFpn. While the carboxylate at the C-terminus of hepcidin participates in the coordination of the bound metal ion to enhance hepcidin affinity, Cys7 of PR73 forms a disulfide bond with Cys326 of Fpn. The density map is of sufficient quality to resolve disulfide bridges, as indicated by the known disulfide bridges in the 11F9 Fab (**Figure 4b** and **Figure 4—figure supplement 2**). Additionally, Tyr333 of TM7b, which forms a hydrogen bond with Met21 of hepcidin, now has a potential cation-π interaction with Arg8 of PR73 (**Figure 4c**). Lastly, the aminohexanoic linker and palmitic amide tail of PR73 protrude out of the domain interface and insert into the membrane. This leads to a novel interaction where Gln194 on TM5 forms a hydrogen bond with the iminodiacetic palmitic amide (Ida(NHPal)) (**Figure 4c**). This interaction is also unique to PR73 as there is no evidence that Gln194 is involved in the hepcidin binding (Billesbølle et al., 2020).

### Validation of PR73-Fpn interactions

PR73 is known to inhibit Fpn in the nanomolar range in cell-based assays (Fung et al., 2015). We measured the binding affinity of PR73 to the purified HsFpn and found that the equilibrium *K*_*D*_ is ∼37 nM (**Figure 5a** and **Table 2**), which is consistent with results from cell-based assays. We then measured binding affinity in the presence of β-mercaptoethanol (BME), which would mask cysteine residues to prevent the formation of disulfide bridges (**Figure 5b** and **Table 2**). The affinity (*K*_*D*_) is reduced by ∼40-fold to *K*_*D*_ ∼1.4 µM in the presence of BME, indicating that the cysteine residues contribute significantly to the binding affinity. We also measured the binding affinity of PR73 and wild-type (WT) HsFpn in the presence of 5 mM CoCl_2_ and that of the Cys326Ser mutant (**Figure 5b**). The Co^2+^ ion was shown to be required for the binding of hepcidin because the C-terminus carboxylate of hepcidin coordinates the ion with Cys326 (Billesbølle et al., 2020). In contrast, PR73 affinity is reduced in the presence of Co^2+^ (*K*_*D*_ ∼880 nM), which we interpret as in the presence of Co^2+^, Cys326 is less available to form a disulfide bridge with PR73. Consistent with this interpretation, the Cys326Ser mutation significantly reduces PR73 affinity (*K*_*D*_ ∼2.4 µM). Although a disulfide bridge between PR73 and HsFpn is resolved in the structure, we observed a measurable off-rate during the dissociation step of the binding assay. This suggests that the disulfide bond formation may not be complete within the duration of the assay or is reversible.

**Figure 5.**
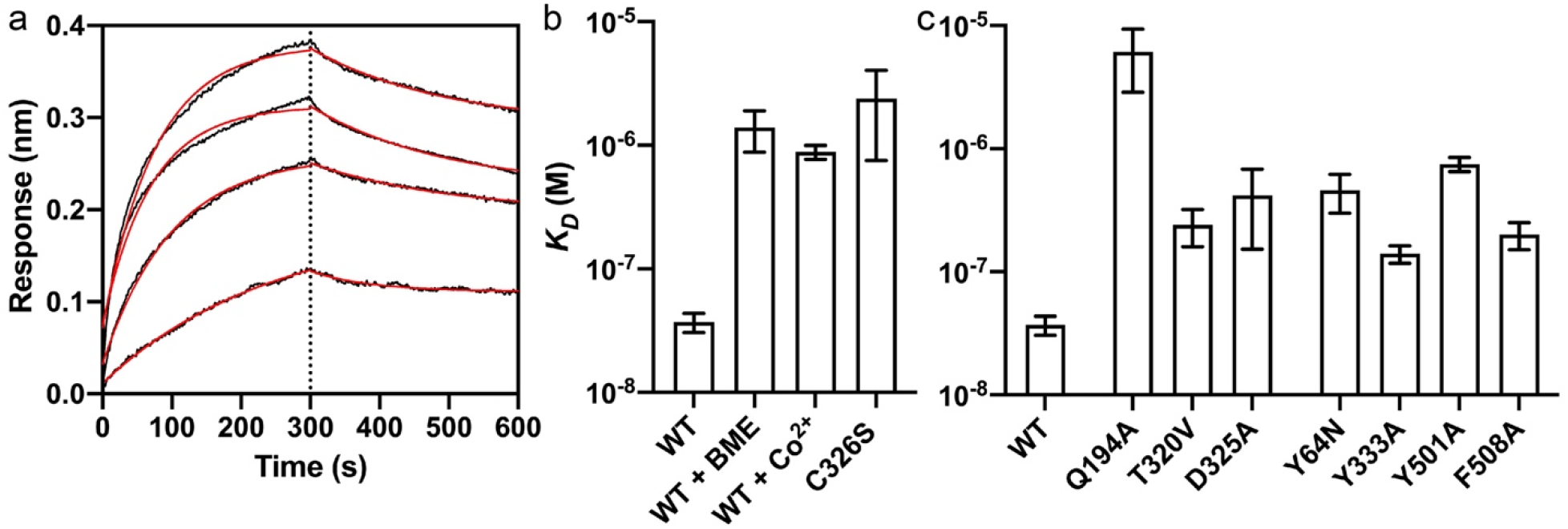
Binding of PR73 to Fpn. **a**. Octet traces of wildtype Fpn in the presence of 1.25, 2.5, 5, and 10 µM PR73. The average *K*_*D*_ is ∼37 nM. **b**. Binding affinity of HsFpn to PR73 in conditions that affect disulfide bridge formation: in the presence of 5 mM BME; 5 mM Co^2+^; Cys326Ser mutation. **c**. Binding affinity of HsFpn mutants to PR73. In this article, the height of the bar graphs represents the mean of at least three measurements and the error bar standard error of the means (SEM).

**Table 1.** Summary of cryo-EM data collection, processing, and refinement.

| Sample | HsFpn-Co<sup>2+</sup>-11F9 | HsFpn-PR73-11F9 |
| --- | --- | --- |
| **Cryo-EM Data Collection** |  |  |
| Voltage (kV) | 300 | 300 |
| Magnification (x) | 81,000 | 81,000 |
| Pixel Size (Å) | 1.08 | 1.10 |
| Electron exposure (e<sup>-</sup>/Å<sup>2</sup>/frame) | 1.25 | 1.25 |
| Defocus range (μm) | [-2.25, -1.0] | [-2.0, -0.8] |
| Number of image stacks | 4,251 | 4,941 |
| Number of frames per stack | 40 | 40 |
| **Cryo-EM Data Processing** |  |  |
| Initial number of particles | 2,175,353 | 2,960,056 |
| Final number of particles | 215,164 | 162,586 |
| Symmetry imposed | C1 | C1 |
| Map resolution (Å) | 3.0 | 2.7 |
| Map resolution range (Å) | 2.5 – 3.7 | 2.3 – 3.8 |
| FSC threshold | 0.143 | 0.143 |
| **Model Refinement** |  |  |
| Number of amino acids | 877 | 905 |
| Total non-hydrogen atoms | 6,185 | 6,569 |
| Average B factor (Å<sup>2</sup>) | 155.5 | 122.76 |
| Bond length RMSD (Å) | 0.004 | 0.003 |
| Bond angle RMSD (°) | 0.817 | 0.782 |
| Ramachandran Plot |  |  |
| Favored (%) | 95.16 | 95.96 |
| Allowed (%) | 4.84 | 3.71 |
| Outliers (%) | 0.00 | 0.34 |
| Rotamer outliers (%) | 2.28 | 1.90 |
| MolProbity Score | 2.30 | 1.91 |

**Table 2.** Binding affinities of PR73 to Fpn measured by Octet BLI. Related to Figure 5.

| Interaction type | Sample name | $K_D$ (M) |
| --- | --- | --- |
| Disulfide bridge by C326 | WT | $3.70 \pm 0.66 \times 10^{-8}$ |
| | WT + BME | $1.39 \pm 0.51 \times 10^{-6}$ |
| | WT + Co<sup>2+</sup> | $8.83 \pm 1.12 \times 10^{-7}$ |
| | C326S | $2.39 \pm 1.64 \times 10^{-6}$ |
| Hydrophilic interactions | Q194A | $6.11 \pm 3.24 \times 10^{-6}$ |
| | T320V | $2.40 \pm 0.80 \times 10^{-7}$ |
| | D325A | $4.18 \pm 2.65 \times 10^{-7}$ |
| Hydrophobic interactions | Y64N | $4.59 \pm 1.60 \times 10^{-7}$ |
| | Y333A | $1.40 \pm 0.23 \times 10^{-7}$ |
| | Y501A | $7.49 \pm 0.98 \times 10^{-7}$ |
| | F508A | $2.01 \pm 0.50 \times 10^{-7}$ |

As further validation of the HsFpn-PR73 structure, we next examined contributions from other binding sites residues. We made alanine mutations to Gln194, Asp325, Cys326, Tyr333, Thr501, and Phe508; and two disease-related mutations Tyr64Asn and Thr320Val. All the mutants have reduced binding affinity. Notably, Gln194Ala has the greatest effect with *K*_*D*_ ∼6.11 µM (**Figure 5c** and **Table 2**), indicating that this novel interaction between Gln194 and Ida(NHPal)11 (**Figure 3c**) plays a significant role in the high potency of PR73.

We then measured inhibition of ion transport by PR73 in a cell-based transport assay. HsFpn was expressed in human embryonic kidney (HEK) cells (**Figure 6—figure supplement 1**), and the cells were loaded with a pH-sensitive fluorescent dye pHrodo Red (**Methods**). The uptake of Co^2+^ by Fpn is accompanied by export of H^+^ (Pan et al., 2020), which leads to a decrease in pH inside the cells. PR73 (2 µM) significantly reduces the transport activity as shown in **Figure 6a**– **b**. Thus, PR73 is confirmed to be a potent inhibitor against Co^2+^ transport by Fpn in HEK cells.

**Figure 6.**
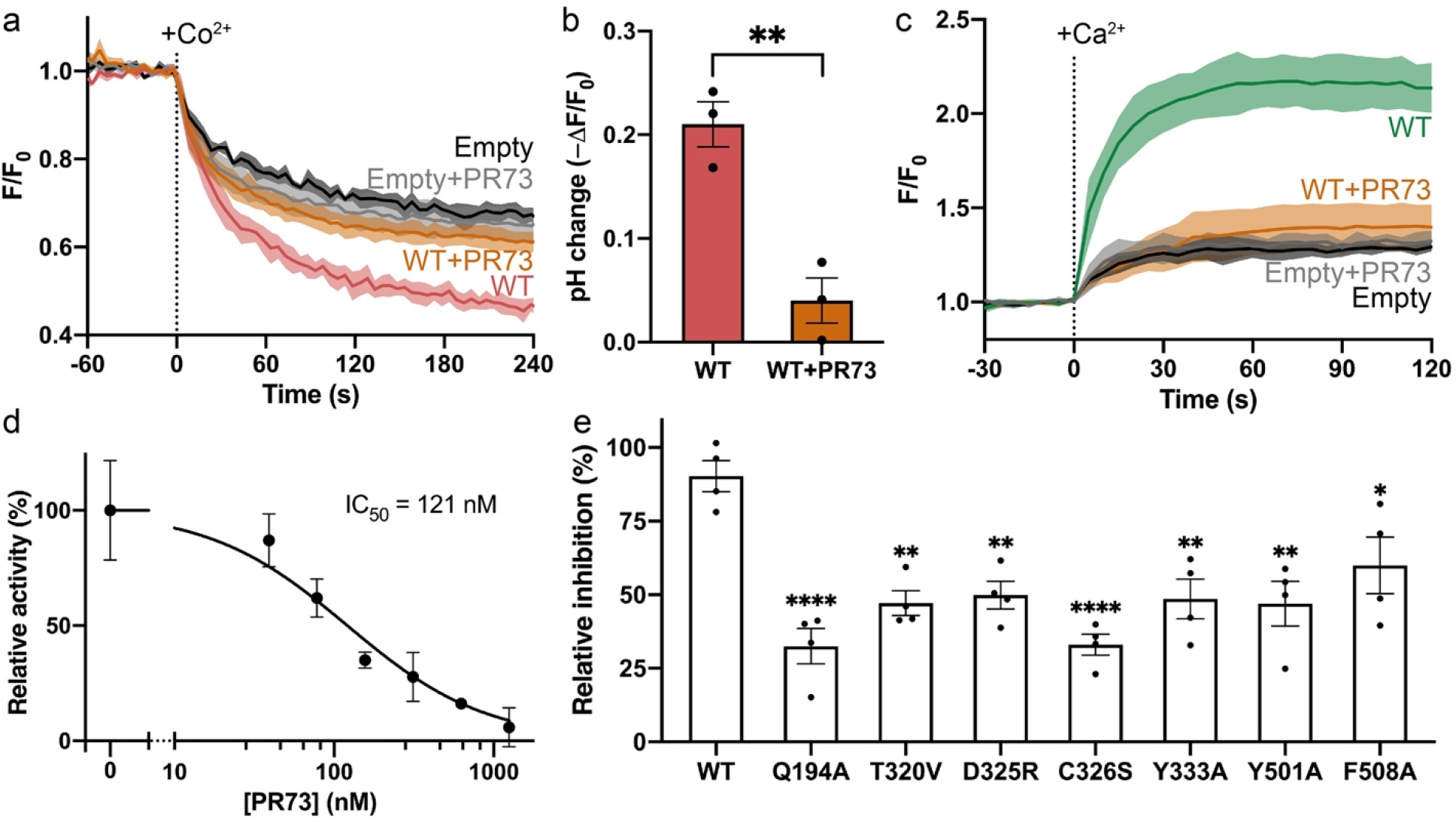
Inhibition of PR73 against the Co^2+^ and Ca^2+^ transport by Fpn. **a**. Co^2+^ import into HEK cells indicated by fluorescence change (F/F_0_) of a pH-sensitive dye (pHrodo Red) loaded inside cells. 500 µM of Co^2+^ was administered at time zero. **b**. Total pH change measured by fluorescence for cells overexpressing Fpn. Data is normalized to empty vector control. The cells overexpressing Fpn show a significantly faster Co^2+^ transport rate. The presence of 50 µM Pr73 significantly reduces transport rate. **c**. Ca^2+^ import into HEK cells measured by fluorescence change (F/F_0_) of a Ca^2+^-sensitive dye (Fluo-4) loaded into cells in the presence of 2 µM PR73. 500 µM of Ca^2+^ was administered at time zero. **d**. Initial rates of fluorescence change versus concentrations of PR73 up to 2 µM. Data are plotted as means with error bars representing the SEM. The solid black line is the data fit to a Michaelis-Menten equation. **e**. Relative inhibition of PR73 binding mutants Fpn proteins. Statistical significance is indicated: *, *p* < 0.05; **, *p* < 0.01; ***, *p* < 0.005; ****, *p* < 0.001.

**Figure 7.**
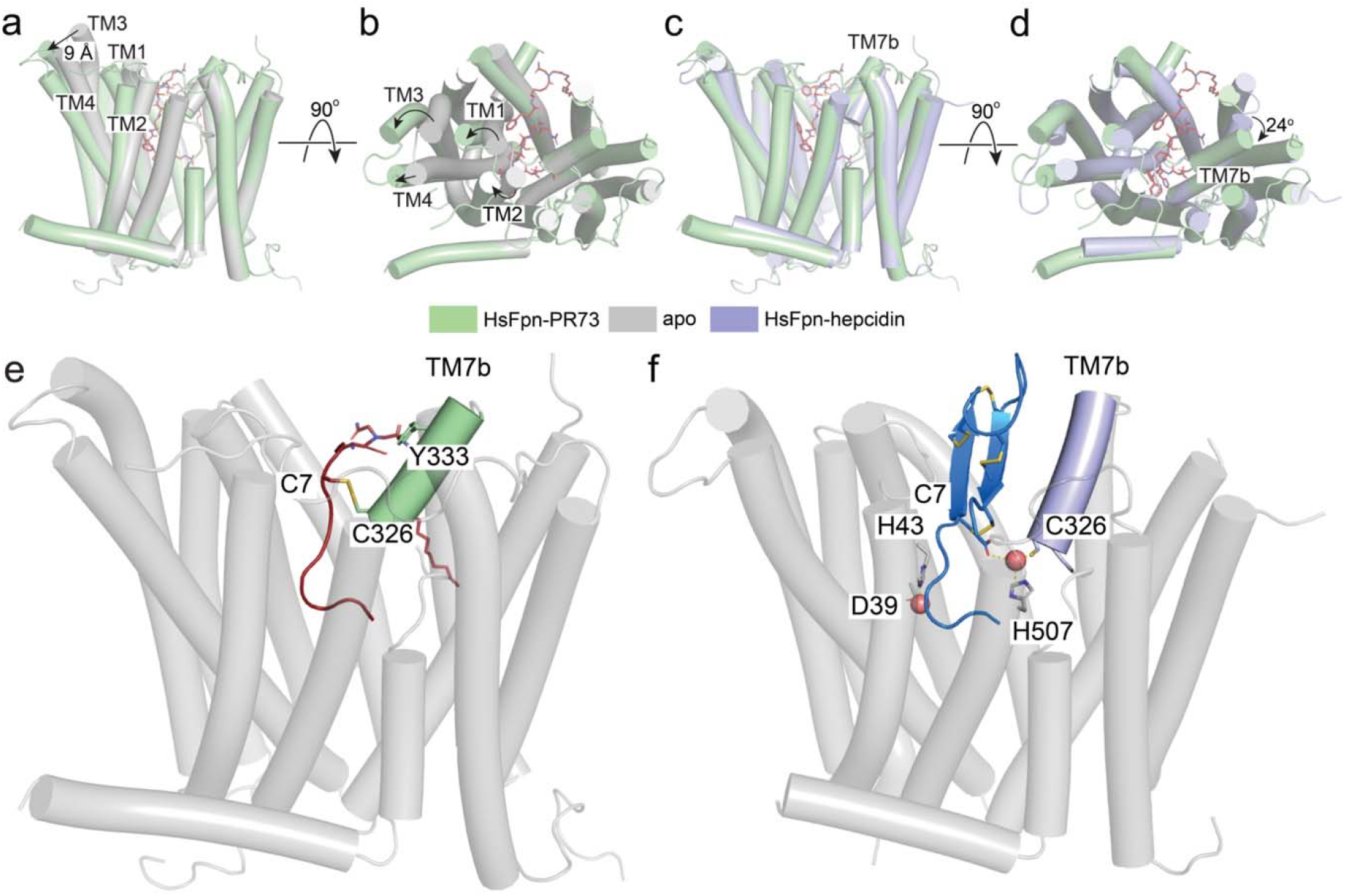
Structural changes induced by PR73. Structure of HsFpn-PR73 (pale green) aligned with apo-HsFpn (grey, PDB ID 6W4S) viewed from the side **(a)** and the top **(b)**. Large structural changes are highlighted by arrows. Structures of HsFpn-PR73 aligned with HsFpn-Hepcidin (light blue, PDB ID 6WIK) viewed from the side **(c)** and top **(d)**. Side views of HsFpn-PR73 **(e)** and HsFpn-Hepcidin (6WIK) **(f)** highlight the disulfide bridge and metal ion coordination. In (**e**), TM7b is colored pale green, and PR73 in brick red. In (**f**), TM7b is colored in light blue, hepcidin shown as marine cartoon with cysteine residues, and C-terminal carboxylate shown as sticks.

We recently showed that Ca^2+^ is transported by HsFpn (manuscript submitted together). We used a Ca^2+^ transport assay to measure the dose-response of PR73 on the activity of HsFpn. PR73 inhibits the Ca^2+^ transport by HsFpn with an IC_50_ ∼121 nM (**Figure 6c–d**). Each of the mutations used in the binding assay was also tested in the cell-based transport assay and exhibited significantly reduced relative inhibition by PR73 (**Figure 6e** and **Figure 6—figure supplement 1**). Results from both the binding and transport assays are consistent with the structure of PR73-bound HsFpn.

## Discussion

Here we have shown two independent inhibition mechanisms of human Fpn. First, we showed that mouse monoclonal 11F9 Fab, which was developed to bind to a monkey homolog of Fpn, binds to HsFpn with nanomolar affinity and inhibits ion transport. The 11F9 interacts with both the NTD and CTD of HsFpn from the intracellular side, and the interactions stabilize the transporter in an outward-facing conformation.

Second, we delineate the inhibition mechanism of PR73 by solving the structure of the minihepcidin in complex with HsFpn. We find that the first six residues of PR73 assume a similar conformation to that of hepcidin, even though the residues are modified to unnatural amino acids, and that these residues interact with HsFpn similarly to hepcidin. However, residues 7 – 11 of PR73 interact with HsFpn differently. While hepcidin has a compact and well-folded structure with four disulfide bridges, PR73 has none. The free Cys7 of PR73 allows for the formation of a disulfide bond to Cys326 of HsFpn (**Figure 7e**). Although it was proposed that hepcidin may form a disulfide bridge with HsFpn, the structure of hepcidin-HsFpn shows that its internal cysteine bridges are intact (**Figure 7f**). We demonstrate that the disulfide bond between PR73 and HsFpn contributes significantly to the binding affinity. The flexibility of PR73 affords another novel interaction between residue 11 of PR73 and Gln194 of HsFpn. This interaction contributes significantly to the binding, and the hydrophobic acyl chain of the palmitic acid may extend into the hydrophobic core of the membrane to further anchor PR73.

Minihepcidins are promising therapeutic reagents being pursued for the treatment of human diseases like β-thalassemia and hemochromatosis (Preza et al., 2011; Ramos et al., 2012). Our study provides a structural framework that highlights unique interactions between PR73 and Fpn and may facilitate and guide future drug development targeting HsFpn.

## Materials and methods

### Cloning, expression, and purification of HsFpn

The cDNA of HsFpn (UniProt ID: <u>Q9NP59</u>) was codon optimized, synthesized, and cloned into a pFastBac dual vector. A Tobacco Etch Virus (TEV) protease site and an octa-histidine (8×His) tag were added to the C-terminus of the protein. The Back-to-Bac method (Invitrogen) was used to express HsFpn was expressed in Sf9 (*Spodoptera frugiperda*). Purification of HsFpn follows the same protocol reported for TsFpn (Pan et al., 2020). Size exclusion chromatography (SEC) was used to collect the purified HsFpn in FPLC buffer consisting of 20 mM HEPES, pH7.5, 150 mM NaCl, and 1 mM (w/v) n-dodecyl-β-D-maltoside (DDM, Anatrace). The Quikchange method (stratagene) was used to generate HsFpn mutations. Mutations were verified by sequencing. Mutant HsFpn proteins were expressed and purified following the same protocol for the WT.

### Octet Biolayer Interferometry

Biolayer interferometry (BLI) assays were performed at 30° C under constant shaking at 1000 rpm using an Octet system (FortéBio). First, amine-reactive second-generation (AR2G) biosensor (Sartorius) tips were activated in 20 mM 1-ethyl-3-[3-dimethylaminopropyl]carbodiimide hydrochloride (EDC) and 10 mM N-hydroxysulfosuccinimide (Sulfo-NHS) for 300 s. Then the tips were immobilized with 5 µg/mL of 11F9 Fab in the FPLC buffer for 600 s. The tips were quenched in 1 M ethanolamine (pH 8.5) for 300 s, The tips with immobilized ligands were equilibrated in the FPLC buffer for 120 s and transferred to wells with a concentration gradient of HsFpn (400, 200, 100, 50, and 25 nM) for 300 s (association) and returned to the equilibration wells for dissociation (300 s). To measure PR73 binding, the tips were immobilized with Fpn at a concentration of 2 µg/mL in the FPLC buffer for 600 s. After quenching the immobilization reaction, the tips were transferred to wells with a concentration gradient of PR73 (10, 5, 2.5, 1.25 µM) for 300 s (association), and back to equilibration wells for 300 s (dissociation). Binding curves were aligned and corrected with the channel of no analyst protein. The association and disassociation phases were fit with 1-exponential functions to extract *k*_*a*_ (association rate constant) and *k*_*d*_ (dissociation rate constant) of the binding, which were used to calculate the dissociation constant *K*_D_.

### Reconstitution of Fpn into liposomes

1-palmitoyl-2-oleoyl-sn-glycero-3-phospho-(1’-rac)-ethanolamine (POPE, Avanti Polar Lipids) and 1-palmitoyl-2-oleoyl-sn-glycero-3-phospho-(1’-rac)-glycerol (POPG, Avanti Polar Lipids) was mixed at 3:1 molar ratio, dried with Argon, and vacuumed overnight to remove chloroform. The lipid was resuspended in reconstitution buffer (20 mM HEPES, pH 7.5, 100 mM NaCl) to a final concentration of 10 mg/mL. The lipid was sonicated until it appeared transparent. 40 mM n-decyl-β-D-maltoside (DM, Anatrace) was added and the sample was incubated for 2 h at room temperature under gentle agitation. HsFpn was added at a 1:100 (w/w, protein:lipid) ratio. Dialysis was performed at 4° C with the reconstitution buffer to remove the detergent. The dialysis buffer was changed daily for three days and then harvested on day 4. Liposome samples were aliquoted, and frozen at -80° C for future use.

### Co^2+^ flux assays in proteoliposomes

Proteoliposome samples of HsFpn were mixed with 250 μM calcein, with or without 20 μM 11F9 Fab, and underwent three cycles of freeze-thaw. The liposomes were extruded to homogeneity with a 400 nm filter (NanoSizer™ Extruder, T&T Scientific Corporation). Excess calcein was removed with a desalting column (PD-10, GE Healthcare) that had been equilibrated with the dialysis buffer. A quartz cuvette was used to detect the fluorescence at 37° C. For the samples loaded with the Fab, an additional 20 μM of 11F9 Fab was incubated with the liposomes prior to reading. The cuvette was read at 10 s intervals with 494 nm excitation and 513 nm emission in a FluoroMax-4 spectrofluorometer (HORIBA). 0.1 mM CoCl_2_ was added to initiate transport.

### Preparation of Fpn-11F9 complex in nanodisc

An established protocol (Martens et al., 2016) was used to express and purify membrane scaffold protein (MSP) 1D1. Lipid preparation was carried out by mixing 1-palmitoyl-2-oleoyl-sn-glycero-3-phospho-(1’-rac)-choline (POPC, Avanti Polar Lipids), POPE, and POPG at a molar ratio of 3:1:1. The lipid mixture was dried with Argon and vacuumed for 2 h. The lipid was resuspended with 14 mM DDM (Autzen et al., 2018). HsFpn, MSP1D1, and the lipid mixture were mixed at a molar ratio of 1:2.5:50 and incubated on ice for 1 h for nanodisc reconstitution. 60 mg/mL of Biobeads SM2 (Bio-Rad) were added three times within 3 h to remove detergents. After the samples were incubated with the Biobeads overnight at 4 °C, the Biobeads were removed. 11F9 Fab was added to the nanodisc sample at a molar ratio of 1.1:1 to Fpn. The complex was incubated on ice for 30 min before it was loaded onto a SEC column that had been equilibrated with 20 mM HEPES, pH 7.5, and 150 mM NaCl. The purified nanodisc sample was concentrated to 10 mg/ml and incubated with 10 mM CoCl_2_ or 1 mM PR73 for 30 min before cryo-EM grid preparation.

### Cryo-EM sample preparation and data collection

The cryo-EM grids were prepared with Thermo Fisher Vitrobot Mark IV. The Quantifoil R1.2/1.3 Cu grids were glow-discharged with air at 10 mA for 15 s using Plasma Cleaner (PELCO EasiGlow™). Aliquots of 3.5 µL of the nanodisc sample were applied to the glow-discharged grids. The grids were blotted with filter paper (Ted Pella, Inc.) for 4.0 s and plunged into liquid ethane cooled with liquid nitrogen. A total of 4,251 (for Co^2+^-bound Fpn) and 4,941 (for PR73-bound Fpn) micrograph stacks were collected on a Titan Krios at 300 kV equipped with a K3 Summit direct electron detector (Gatan) and a Quantum energy filter (Gatan) at a nominal magnification of 81,000 × and defocus values from -2.25 to -1.0 µm (HsFpn-Co^2+^) or - 2.5 to -0.8 µm (HsFpn-PR73). Each stack was exposed for 0.0875 s per frame in the super-resolution mode for a total of 40 frames per stack, which results in a total dose of ∼50 e^-^/Å^2^. The stacks were motion corrected with MotionCor2 (Zheng et al., 2017) and binned 2-fold. The final pixel size is 1.08 Å/pixel (HsFpn-Co^2+^) or 1.10 Å/pixel (HsFpn-Pr73). In the meantime, dose weighting was performed (Grant & Grigorieff, 2015). The defocus values were estimated with Gctf (Zhang, 2016).

### Cryo-EM data processing

A total of 2,175,353 (for Co^2+^-bound Fpn) and 2,960,056 (for PR73-bound Fpn) particles were automatically picked based on a reference map of TsFpn-11F9 (EMD-21460) that was low-pass filtered to 20 Å in RELION 3.1 (Kimanius et al., 2016; Scheres, 2012; Zivanov et al., 2018). Particles were extracted and imported into CryoSparc (Punjani et al., 2017) for 2D classification. a total of 1,242,825 (HsFpn-Co^2+^) and 918,469 particles (HsFpn-PR73) were selected from good classes in 2D classification. 100,000 particles for the good classes were used to generate four initial reference models. Multiple rounds of heterogeneous refinement were performed with particles selected from the 2D classification until < 5% input particles were classified into bad classes. A total of 215,164 particles (HsFpn-Co^2+^) or 454,601 particles (HsFpn-PR73) were subjected to non-uniform (NU) refinement. After handedness correction, local refinement and CTF refinement were performed, resulting in a reconstruction with an overall resolution of 3.0 Å for HsFpn-Co^2+^ and 2.6 Å for HsFpn-PR73. Additional rounds of heterogenous refinement were performed for HsFpn-PR73 with three reference models of “class similarity” of 1 to further improve the density for PR73. Classes with strong densities for PR73 were selected and subjected to NU refinement. The final total of 162,586 particles yielded a reconstruction with an overall resolution of 2.7 Å for HsFpn-PR73. Resolution was estimated with the gold-standard Fourier shell correlation 0.143 criterion (Rosenthal & Henderson, 2003). Local resolution of the maps was estimated in CryoSparc (Punjani et al., 2017).

### Model building and refinement

The structure of apo HsFpn (PDB ID 6W4S) and the 11F9 Fab (from PDB ID 6VYH) were individually docked into density maps in Chimera (Pettersen et al., 2004). The docked model was manually adjusted in COOT (Emsley et al., 2010). PHENIX (Adams et al., 2010) was used for real space refinements with secondary structure and geometry restraints. The EMRinger Score (Barad et al., 2015) was calculated for the models. Structure figures were prepared in Pymol and ChimeraX (Pettersen et al., 2021).

### Ca^2+^ influx and H^+^ transport assays in HEK cells

The pEG BacMam plasmids with WT or mutant Fpn or the empty plasmid were transfected into HEK 293S cells on black wall 96-well microplates coated with poly-D-lysine (Invitrogen/Thermo Fisher). After 2 days, cells were washed in the live cell imaging solution (LCIS) containing 20 mM HEPES, 140 mM NaCl, 2.5 mM KCl, 1.0 mM MgCl_2_, 5 mM D-glucose, pH 7.4. The loading of Fluo-4 (Invitrogen/Thermo Fisher, AM, cell-permeant) for Ca^2+^ uptake or pHrodo™ Red (Invitrogen/Thermo Fisher, AM) for H^+^ transport was performed following the manufacturer’s protocols. After dye loading finished, free dyes were washed away, and cells in each well were maintained in 90 µL LCIS. Both the Ca^2+^ uptake and H^+^ transport was performed in the FlexStation 3 Multi-Mode Microplate Reader (Molecular Devices) at 37 LJ. Fluorescence changes were recorded at an excitation and emission wavelength of 485 nm and 538 nm for Ca^2+^ uptake, and 544 nm and 590 nm for H^+^ transport with 5 s intervals. For PR73 inhibition, cells were incubated with desired concentrations of PR73 for 5 mins prior to reading. Transport was initiated by the addition of 10 µL ligand stock solution (CaCl_2_ or CoCl_2_) to achieve 500 µM extracellular Ca^2+^ or Co^2+^. For the Ca^2+^ uptake, the slopes of straight lines fitted to transport data within 25 s were used to represent initial rates. For the H^+^ transport, relative fluorescence changes at the equilibrium stage were averaged to represent intracellular pH changes.

## Data availability

The cryo-EM density maps of nanodisc-encircled HsFpn-11F9 bound to PR73 or Co^2+^ have been deposited in the Electron Microscopy Data Bank (https://www.ebi.ac.uk/pdbe/emdb/) under accession codes EMD-27498 and EMD-27499, respectively. The corresponding atomic coordinate files have been deposited in the Protein Data Bank (http://www.rcsb.org) under ID codes 8DL7 (HsFpn-PR73) and 8DL8 (HsFpn-Co^2+^). Any additional information required to reanalyze the data reported in this paper is available from the lead contact Yaping Pan upon request.

## Acknowledgments

This work was supported by grants from the NIH (HL157473 to Y.P. and DK122784 and HL086392 to M.Z.), and Cancer Prevention and Research Institute of Texas (R1223 to M.Z.). We acknowledge the use of the cryo-EM core at Baylor College of Medicine (BCM) for grid preparation and screening. Cryo-EM data in this work were acquired at the Pacific Northwest Center for Cryo-EM (PNCC) at Oregon Health & Science University, supported by the NIH grant U24GM129547, and the National Center for Cryo-EM Access and Training (NCCAT) and the Simons Electron Microscopy Center at the New York Structural Biology Center, supported by the NIH grant U24 GM129539 and by grants from the Simons Foundation (SF349247) and NY State Assembly. We acknowledge L. Wang for help with grid preparation, and Z. Ren for making some of the mutations. We are grateful to A. A. R. Adeosun for her instructions on the use of FlexStation 3.

## Author Contributions

Y.P., M.Z., A.S.W. and J.S. conceived the project. J.S., A.S.W. and Y.P. conducted experiments. P.R. provided PR73. Y.P., M.Z., A.W.S., and J.S. wrote the manuscript. All authors contributed to the revision of the manuscript.

## Competing interests

The authors declare no competing financial interests.

---

**Figure 2—figure supplement 1.**
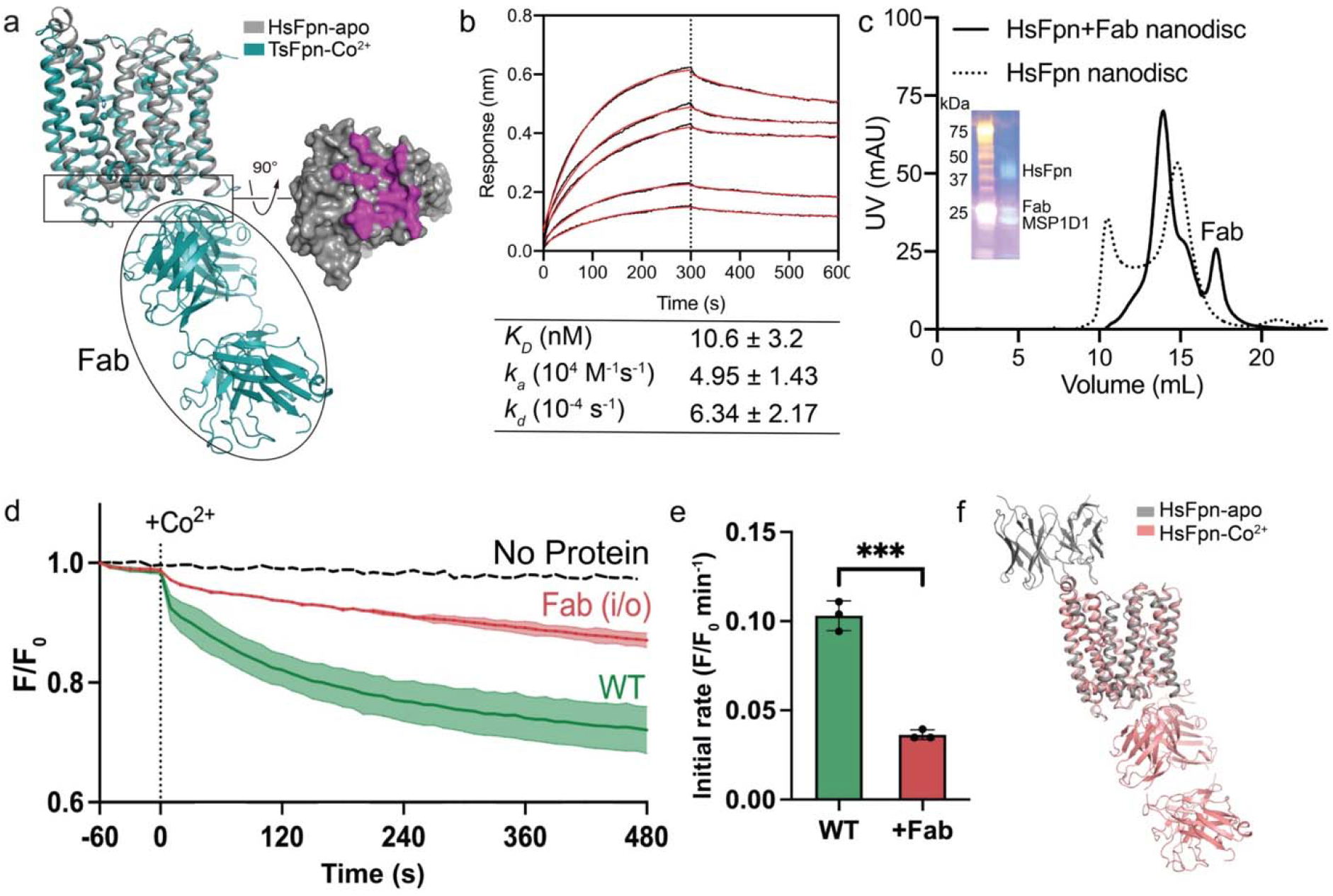
Purification and reconstitution of HsFpn-11F9 complex. **a**. Left: structural alignment of HsFpn (grey, PDB ID 6W4S) and TsFpn (teal, PDB ID 6VYH). Right: the intercellular side of TsFpn is shown as a grey surface with the epitope of 11F9, defined as residues within 4 Å from 11F9, colored in magenta. **b**. Binding of 11F9 Fab to HsFpn measured by Octet BLI. **c**. Size-exclusion chromatography (SEC) profiles and SDS-PAGE gel image (inset) of HsFpn-11F9 in nanodisc. HsFpn-11F9 complex (solid line) eluted significantly earlier than HsFpn alone (dotted line). The later peak from the complex sample comes from the Fab in excess. **d**. Co^2+^ import into proteoliposomes measured by fluorescent changes of (F/F_0_) of a transition metal ion-sensitive dye (calcein). Addition of the Fab to the inside and outside of the liposome inhibits HsFpn transport activity. 100 µM Co^2+^ was added at time zero. **e**. Initial rates of fluorescence change with and without Fab added. The height represents the mean of at least three measurements and the error bar standard error of the means (SEM). Student’s t-test *p* = 0.0013. **f**. Structural comparison between apo (PDB ID 6W4S) and Co^2+^-bound HsFpn. Notice that the Fab, used by Billesbølle et al. for structural determination, binds to the extracellular side and interacts only with the NTD of Fpn.

**Figure 2—figure supplement 2.**
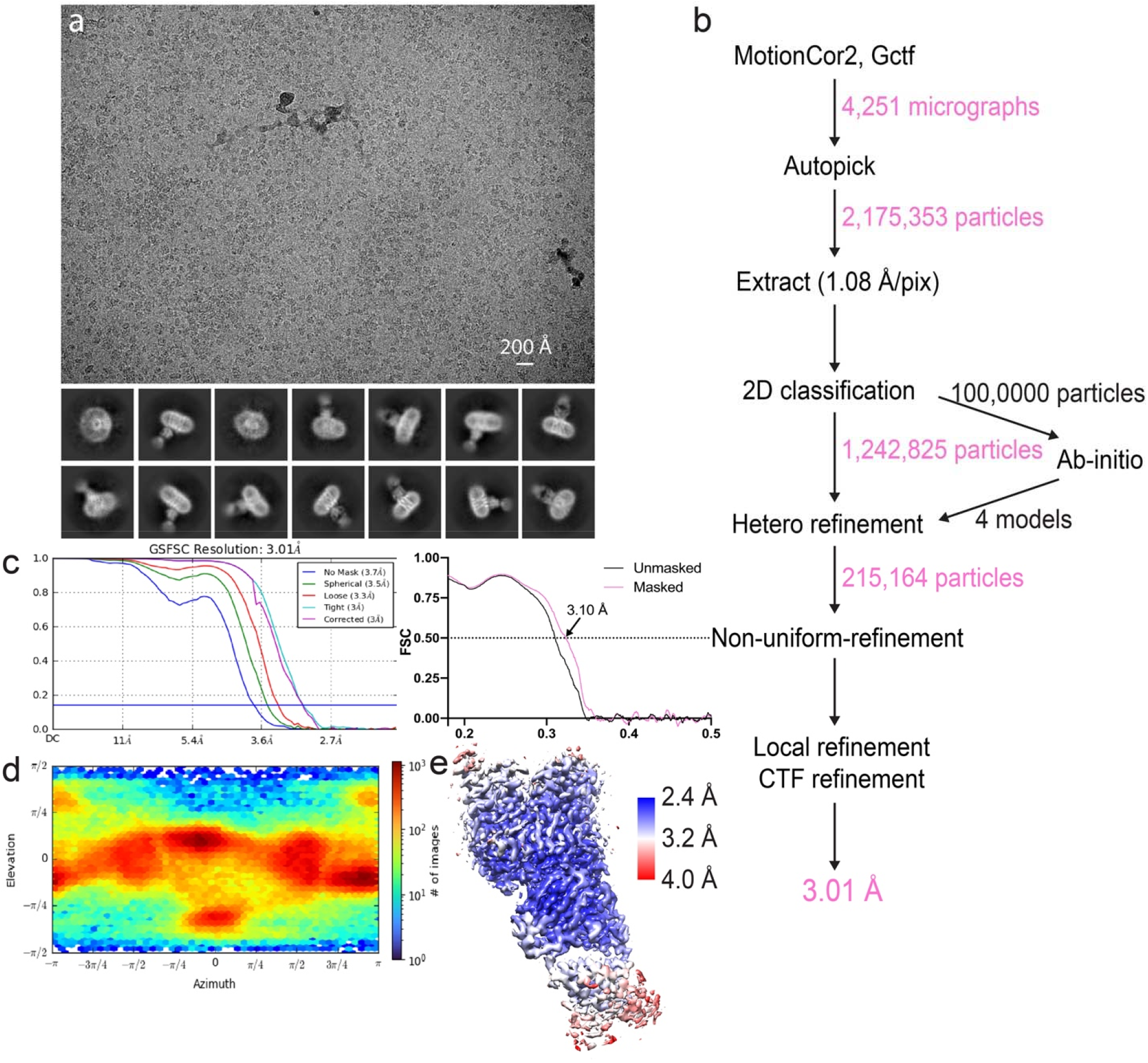
Cryo-EM analysis of HsFpn-Co^2+^ in nanodisc. **a**. Representative electron micrograph (upper panel) and 2D class averages (lower panel). **b**. Workflow of data processing for single-particle reconstruction. **c**. The gold-standard Fourier shell correlation (FSC) curves for the final map (left panel) and map-to-model FSC curves (right panel). **d**. Direction distribution of particles used in the final 3D reconstruction. **e**. Local resolution map colored from 2.4 Å (blue) to >4.0 Å (red).

**Figure 2—figure supplement 3.**
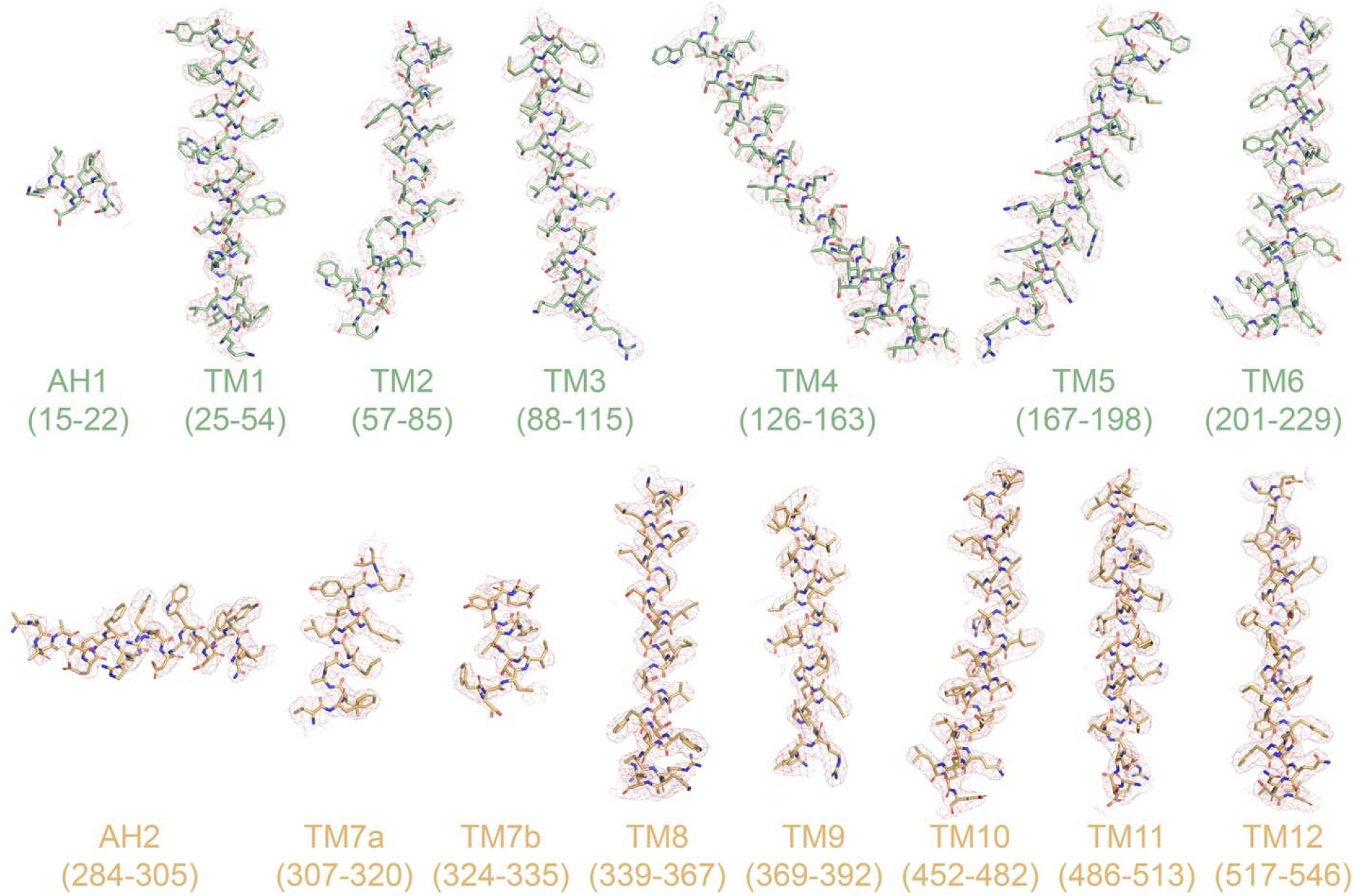
Cryo-EM densities of TM helices and amphipathic helices (AH). Densities for HsFpn-Co^2+^ are shown as pink mesh. Residues within the ranges indicated below are rendered in stick representations and colored in pale green for NTD and light orange for CTD.

**Figure 3—figure supplement 1.**
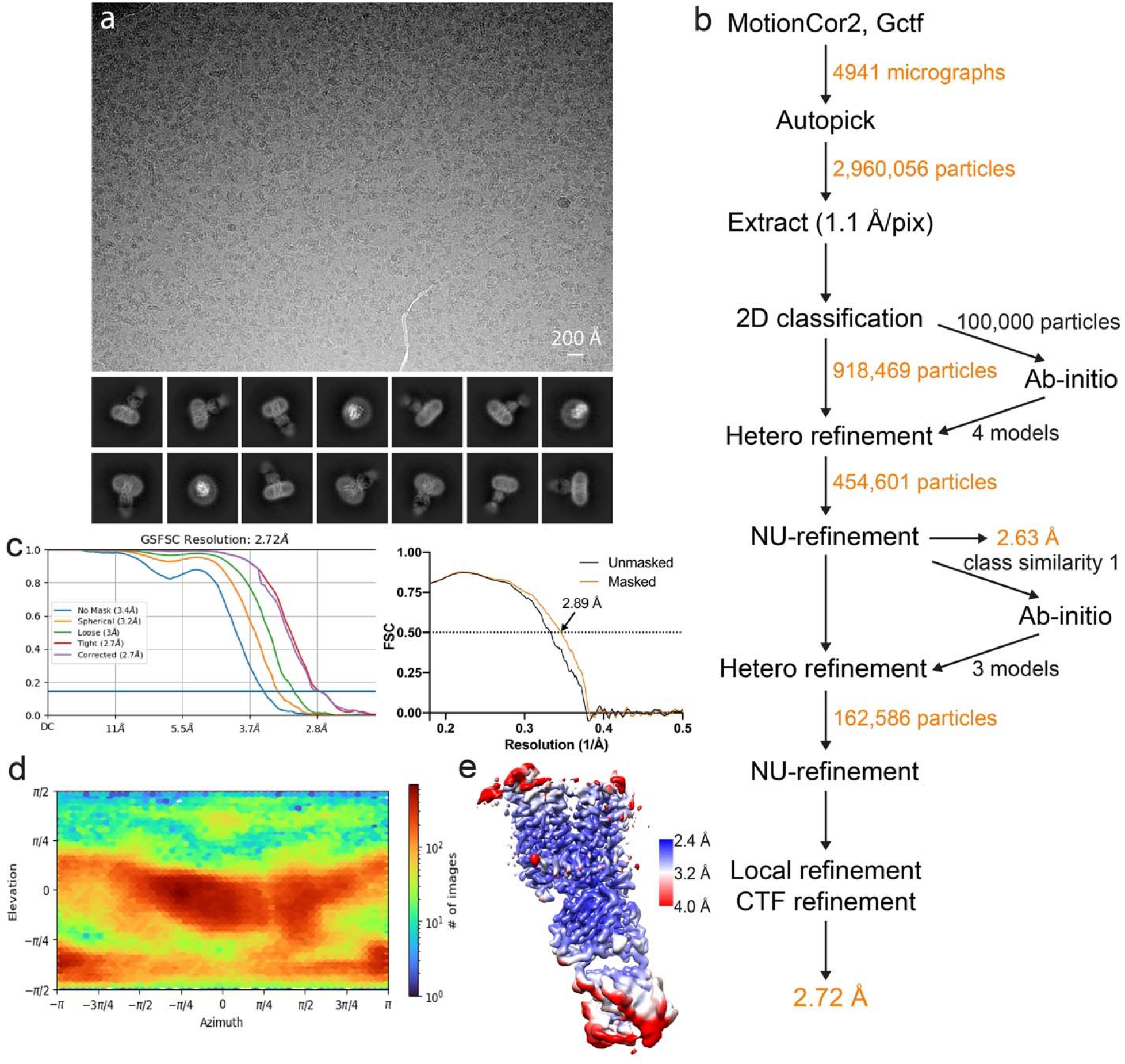
Cryo-EM analysis of HsFpn-PR73 in nanodisc. **a**. Representative electron micrograph (upper panel) and 2D class averages (lower panel). **b**. Workflow of data processing for single-particle reconstruction. **c**. The gold-standard Fourier shell correlation (FSC) curves for the final map (left panel) and map-to-model FSC curves (right panel). **d**. Direction distribution of particles used in the final 3D reconstruction. **e**. Local resolution map colored from 2.4 Å (blue) to >4.0 Å (red).

**Figure 3—figure supplement 2.**
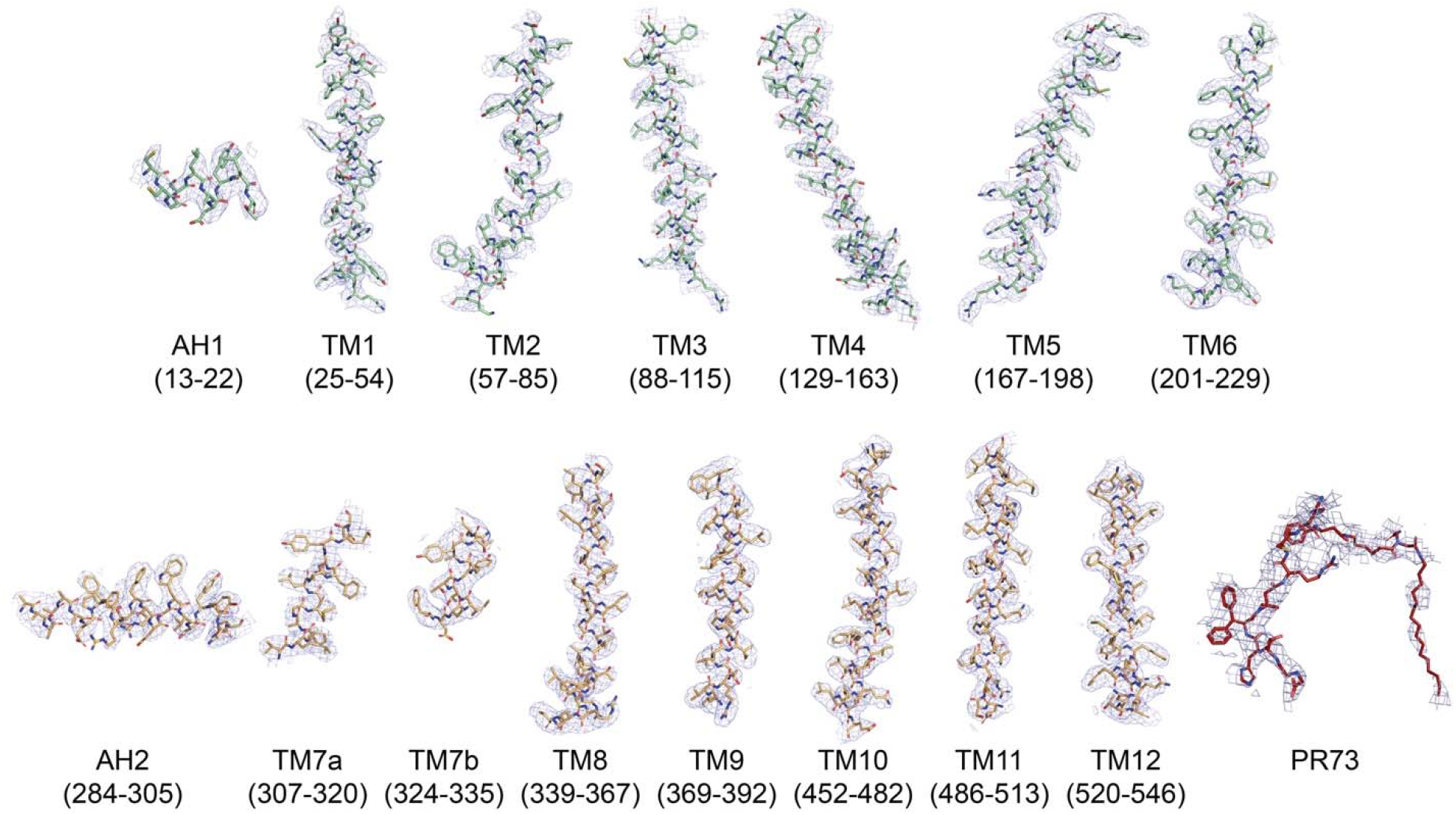
Cryo-EM densities of transmembrane (TM) helices and amphipathic helices (AH) in HsFpn-PR73. Densities of HsFpn-PR73 are shown as blue mesh. Residues within the ranges indicated below are represented as sticks and colored in pale green, light orange, or brick red for the NTD, CTD, and PR73 respectively.

**Figure 3—figure supplement 3.**
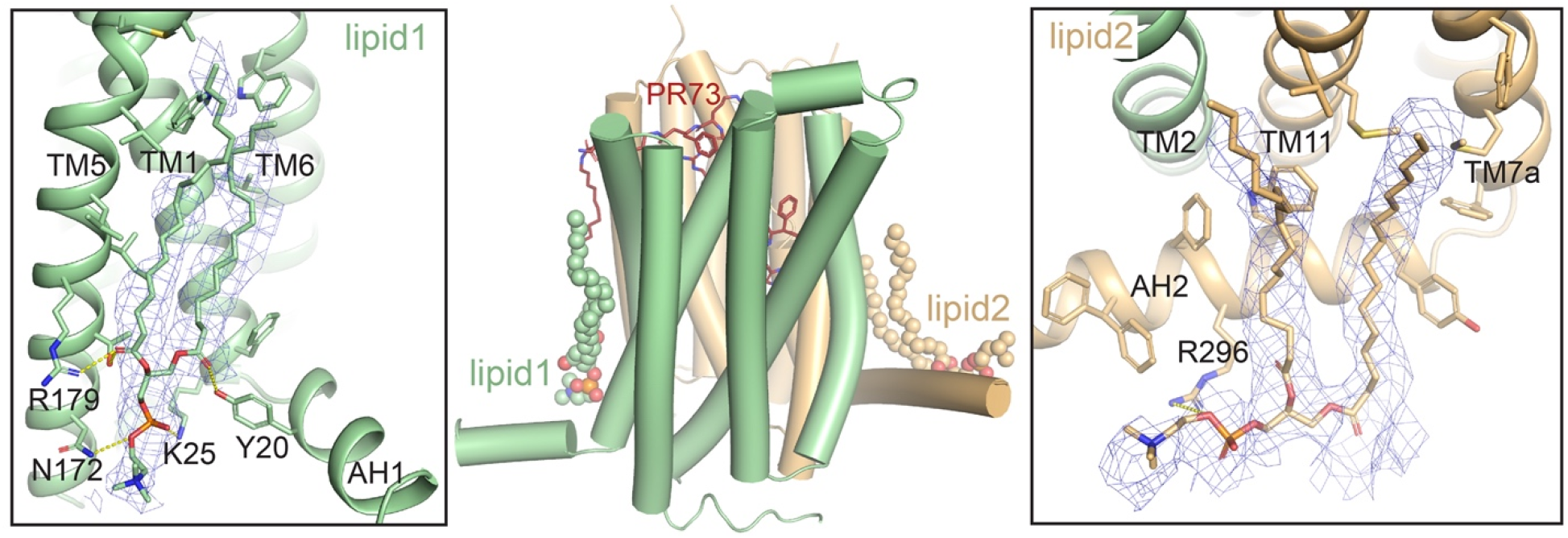
Lipid molecules bound to HsFpn. Side view of two phospholipids (shown as spheres) located on the intracellular side in the HsFpn-PR73 structure (middle panel). Interactions of these lipids with residues of HsFpn (left and right panels). Side chains of residues within 4 Å from the lipids are shown as sticks. Hydrophilic interactions are indicated with yellow dashed lines. Densities of the lipids are contoured at 3.5σ as blue mesh.

**Figure 4—figure supplement 1.**
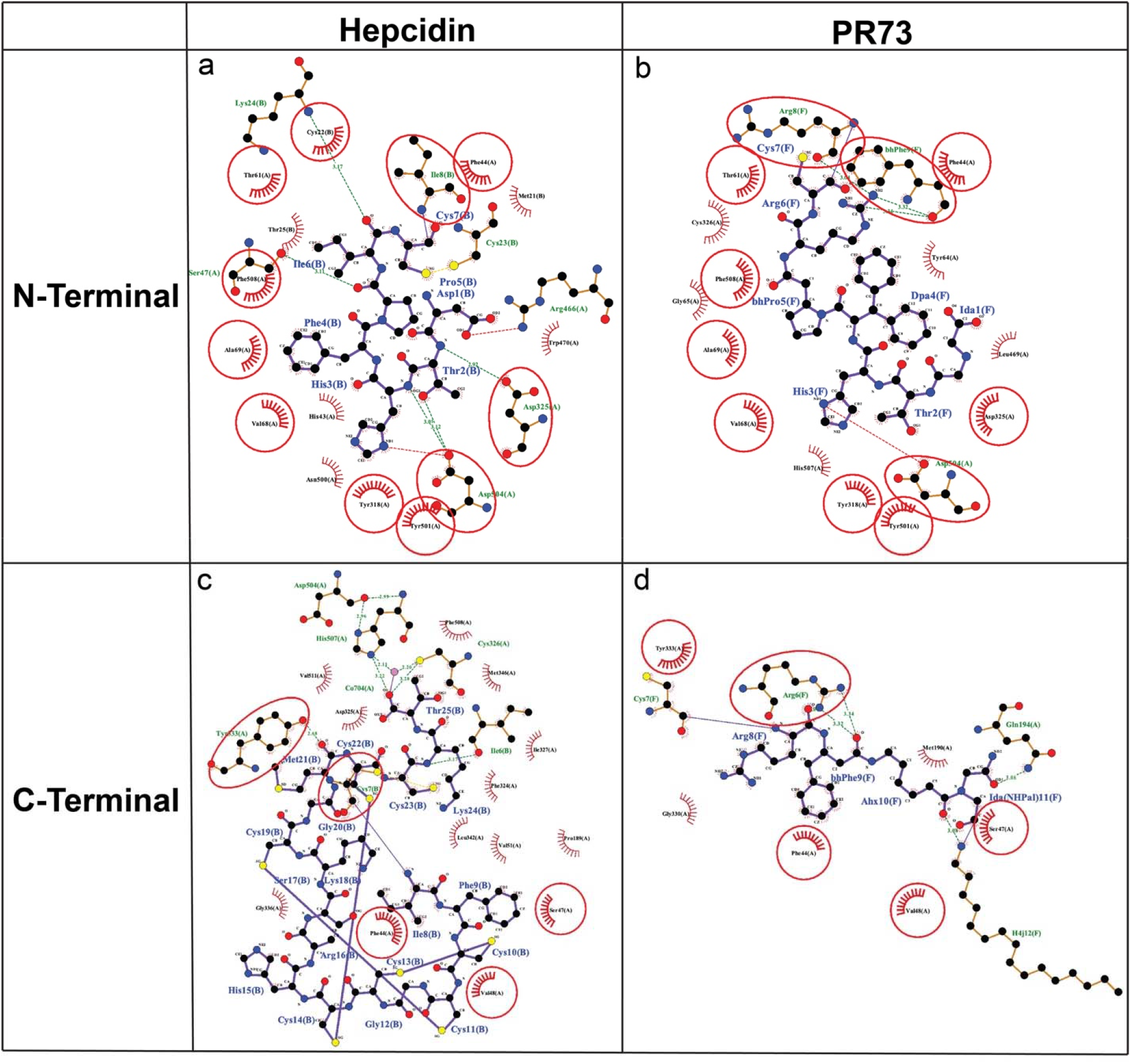
Binding site interactions PR73 vs hepcidin. Ligplot of hepcidin **(a, c)** and PR73 **(b, d)** showing interactions in the binding pocket of HsFpn. The peptide is shown as purple sticks. Interactions common in both structures are marked by red circles.

**Figure 4—figure supplement 2.**
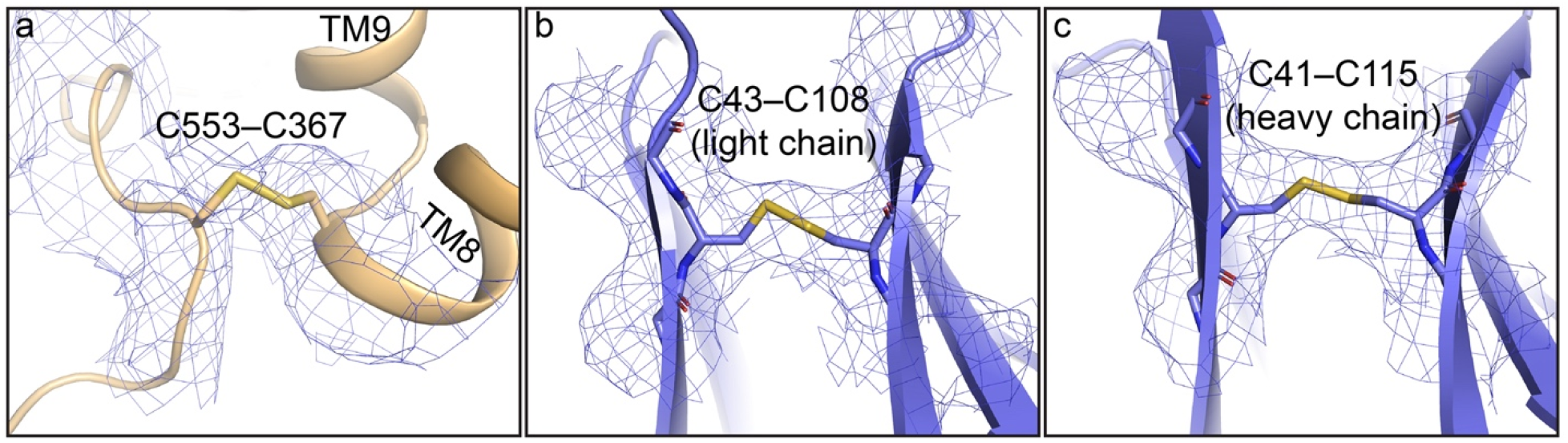
Native disulfide bridges in the HsFpn-PR73 Fab structure. **a**. Cys553 near the C-terminus of HsFpn forms a disulfide bond with Cys367 between TM8 and TM9. Disulfide bonds in the light chain (**b**) and heavy chain (**c**) of the Fab. All density maps are contoured at 4.5σ as blue mesh.

**Figure 6—figure supplement 1.**
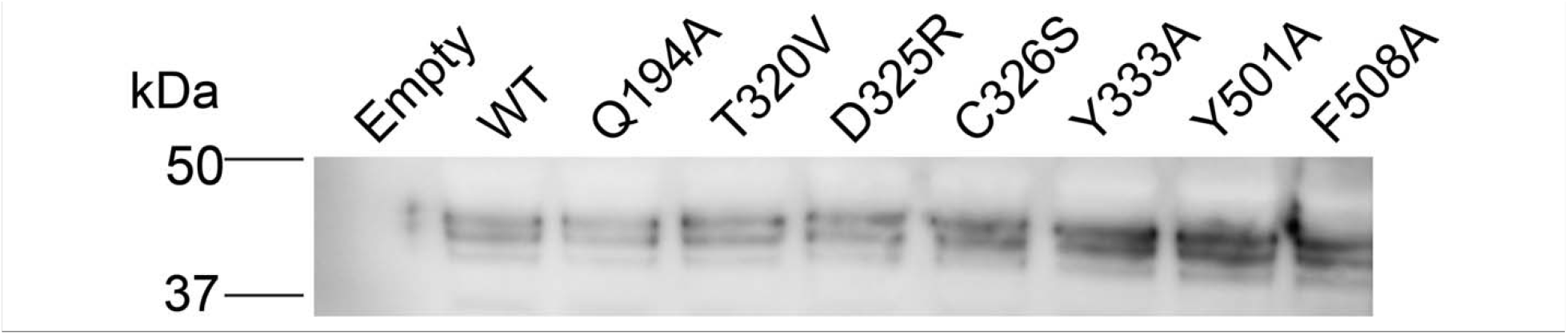
Expression of WT and mutant Fpn in HEK cells assessed by western blot.

## Notes

### Competing Interest Statement

The authors have declared no competing interest.

